# Metformin an anti-diabetic drug, possess ACE-2 receptor-SARS-Cov-2 RBD binding antagonist activity, anti-inflammatory and cytokine inhibitory properties suitable for treatment of COVID-19

**DOI:** 10.1101/2023.05.14.540726

**Authors:** Uday Saxena, Kranti Meher, K Saranya, Subrahmanyam Vangala

## Abstract

Metformin is a widely used and is a safe anti-diabetic drug. It has also been shown to have anti-inflammatory and anti-viral activities in humans and animal models. Specifically we explored its activity in SARS-CoV-2 initiated COVID19 disease. Here we show that metformin 1. blocks the binding of SARS-CoV-2 spike protein receptor binding domain RBD to human ACE2 receptor 2. We also show that it has anti-inflammatory effects and reduces cytokine secretion as well as blocks the recruitment of monocytes to endothelial cells 3. Finally we show its activity in a hamster in vivo model of SARS-CoV-2 infection as a nasal formulation. Based on the safety and the therapeutic properties relevant to COVID-19 it is feasible to propose a nasal spray of metformin that can be used in treatment of this disease. A nasal spray would deliver the drug to the target organ lung and spare other organs which get exposed upon oral dosing.

## Introduction

The progression of COVID-19 disease starts with the SARS-CoV-2 virus attaches itself to human host cells such as nasal epithelial and lung cells. 1. It binds to ACE2 receptor on these cells using the receptor binding domain (RBD) on its surface spike protein. The three steps that allow infection are: 1. Once bound it penetrates into the cells and starts to replicate. 2 It simultaneously creates an inflammatory environment and leukocytes are recruited to the lungs by the binding of these cells to lung endothelium. 3. These leukocytes can further extend the inflammation by massive production of cytokines which is called cytokine storm. This worsens the disease and ultimately the lungs become dysfunctional (1,2).

Repurposed drugs represent an accelerated way of bringing therapies to patients. A repurposed drug is typically a drug which has been approved for an indication and can be used for a new indication. This approach saves time and money because the drug has undergone several safety and regulatory requirements prior to market launch and thus the work needed on these fronts is less than a brand new drug.

We hypothesize that a single drug that can attack all three steps mentioned above will be most effective in fighting COVID-19 rather than a drug that only treats one step. We choose metformin as a potential candidate drug because of its known human safety and the demonstrated anti-viral and anti-inflammatory properties drug in literature and can be rapidly repurposed for COVID19 (3). We show robust data that metformin can antagonize the binding of RBD to ACE2, thus can prevent viral entry, can block inflammation cascade initiated by monocyte recruitment and finally it showed ability to block IL-1b secretion from human lung cells. Based on these properties we tested a to a nasal formulation of metformin as treatment for COVID-19 in the hamster model and show activity in vivo.

## Methods

### 1. RBD-ACE2 binding assay

The above assay was performed using SARS-CoV-2 sVNT ready to use kit by Genscript, which is a competition ELISA (4). In the first step, a mixture of horse radish peroxidase-RBD (HRP-RBD) and controls/drugs (drugs at the concentrations of 50μM, 100μM and 200μM) were incubated at 37o C for an hour to allow the binding of drugs to HRP-RBD. Following the incubation, these mixtures were added to a capture plate, which was coated with human ACE-2 receptor to allow binding of any unbound HRP-RBD and the ones bound to drugs to ACE-2 receptor. After incubating the microplate at 37o C for an hour, it was washed four times using wash buffer in order to remove any unbound HRP-RBD. Washing step was followed by addition of colour substrate tetramethyleneblue (TMB), turning the colour to blue. The reaction was allowed to run for 15 minutes followed by stop solution turning the colour from blue to yellow which was read at 450 nm.

### 2. IL-1*beta (IL1b)* secretion assay

HUVEC cells were grown at 37o C in the Endothelial cell basal medium-2 (LONZA #Cat No-CC-3156 and CC-4176) and were used with or without addition of drugs and LPS (0.5 ug/ml) (4). Human IL-1b ELISA was performed to detect the presence of IL-1b secreted in the media. The ELISA was performed using Human IL-1b high sensitivity kit from Invitrogen. The samples for ELISA were media collected after treating the cells with drugs at 100uM. Samples were added to the microplate precoated with human IL-1b and secondary anti-human IL-1b antibody conjugated to biotin was added to the plate.

### 3. Monocyte accumulation assay

Our 3D vascular lung model was used to study the adhesion of monocytes to the endothelium (4,5). This model consists, A549 lung cell line and HUVEC endothelial cells. For monocytes, a human monocytic cell line HL60 cells, were grown at 37o C in the growth medium RPMI (HIMEDIA #Cat No-AT028) supplemented with 10% (v/v) fetal bovine serum under the atmosphere containing 5% CO2.

#### 3D-bioprinting

In the 3D-vascular lung model two cell layers were bioprinted, first A549 cell suspension was loaded into bioprinter syringe and printed in each well of 96 well plate. After 48 hours of incubation, HUVEC cells were printed on top. After 48 hours of incubation the media was removed and they were added with lipopolysaccharide (LPS) at 0.5 μg/ml and without LPS at the desired concentration. Later, cells were incubated overnight. Next day the drugs were removed and the cells were washed once with media. For studying monocyte adhesion to the endothelium, MTT stained HL60 monocytic cells 1x 104 cells were and incubated the plate for and the wells and pictures were taken of all the wells and the bound HL 60 cells were counted with the help of ImageJ, an open access software platform.

### 4. Cytotoxicity and Viral load assay in Vero E6 cells

The cytotoxicity assay of the test compounds for cell death(cytotoxic) of the cells was performed using an LDH release assay in Vero E6 cells. The viral load plaque assay was performed using Vero E6 cells in which SARS-CoV-2 infection causes substantive cell death. These assays were performed Foundation of Neglected Research (FNDR, Bangalore, India) in their BSL3 labs. The 50% inhibitory concentration (IC_50_) is the concentration of the drug which inhibits virus-induced cytotoxic halfway between the baseline and maximum.

### 5. Animal studies

#### NASAL formulation selection

Prior to using a formulation we tested for solubility of metformin in deionised water, saline and CMC tween (microcrystalline cellulose and carboxymethylcellulose sodium, dextrose, 0.02 w/w benzalkonium chloride, polysorbate 80, and 0.25 w/w phenylethyl alcohol, together with metformin). Since metformin was completely soluble in both water and saline we used these formulations rather than CMC Tween, to test the effects of the drug on SARS-CoV-2 infection of Vero E6 Cells at Foundation of Neglected Diseases Research (FNDR, Bangalore, India) a fully accredited lab to perform the Vero E6 cells and the hamster in vivo studies of SARS-Cov-2 infection in their BSL3 safety facility.

#### Hamster STUDY PLAN

Syrian Golden Hamsters (Female) Approximately 10 to 12 weeks of age at the time of intake into study (Average weight ∼ 80 - 100 grams at intake) with 5 animals per group were used.

##### Control groups -

1. Mock Control (non-infected)
2. Infected Vehicle Control (virus and vehicle)
3. Positive Control (virus and remdesivir as a positive control drug)

##### Treatment groups

4. Metformin 10mg/ml
5. Metformin 50mg/ml

Drug dosing: After 7 days acclimatization in BSL3 lab, drug dosing by intranasal route was -1 day dosing pre-infection, then 1 day infection, 3 days of drug dosing, day 4 sacrifice of animals and plating for PFU counts. Metformin in deionized water formulation at 10 mg/ml and 50 mg/ml groups was be given in the morning of Day 0, followed by infection after 6 hours of the dose. This is to reduce the risk associated with giving 100 uL of liquid through intubation twice in the nares on the same day (once for infection and once for test drug).

#### Lung viral load estimation via PFU analysis

After sacrifice at Day 4 post infection for all groups the Standard PFU assay was performed in Vero-E6 cells. The following were recorded:

1. Percentage reduction in viral load in lungs in test material groups vs virus-only group
2. Percentage change in body weight of animals in all groups
3. Lung weight at sacrifice for all animals.

#### Standard PFU assay in Vero-E6 cells to estimate viral load in tissues

Six serial log dilutions of each sample (homogenized tissue extracted in PBS) were added onto pre-formed Vero E6 cell monolayers in a 96-well cell culture plate and incubated for 1 h at 37 °C in 5% CO2 incubator. The samples were then removed. The cell monolayer is then overlaid with cell culture medium containing equal volumes DMEM (2x) and 2% carboxymethylcellulose and incubated for 3 days at 37 °C in 5% CO2, at which time the plates are fixed with formaldehyde and stained with crystal violet.

Plaques are counted for each sample at the dilution at which clear readable counts are possible and the plaque forming units PFU are calculated. Untreated cells alone were used as negative control.

## Results

### 1. Antagonistic activity of metformin towards ACE2-RBD interaction

We first examined whether metformin can block the binding of SARS-Cov-2 RBD protein to ACE2 receptors using a biochemical assay. This biochemical assay addresses the binding interactions without the complexity of cell, metabolism and permeability. We found (Figure 1) that metformin at 100 and 200 uM completed blocked the binding of RBD to ACE2 receptors. Homoharringtonine was another compound that inhibited the activity (we previously reported this activity-reference 5). Several other compounds of diverse chemical structures were tested and showed no inhibitory activity, suggesting selectivity for metformin. This inhibitory activity of metformin suggests that it may prevent the entry of the virus into human cells through the ACE2 receptors.

**Figure 1:**
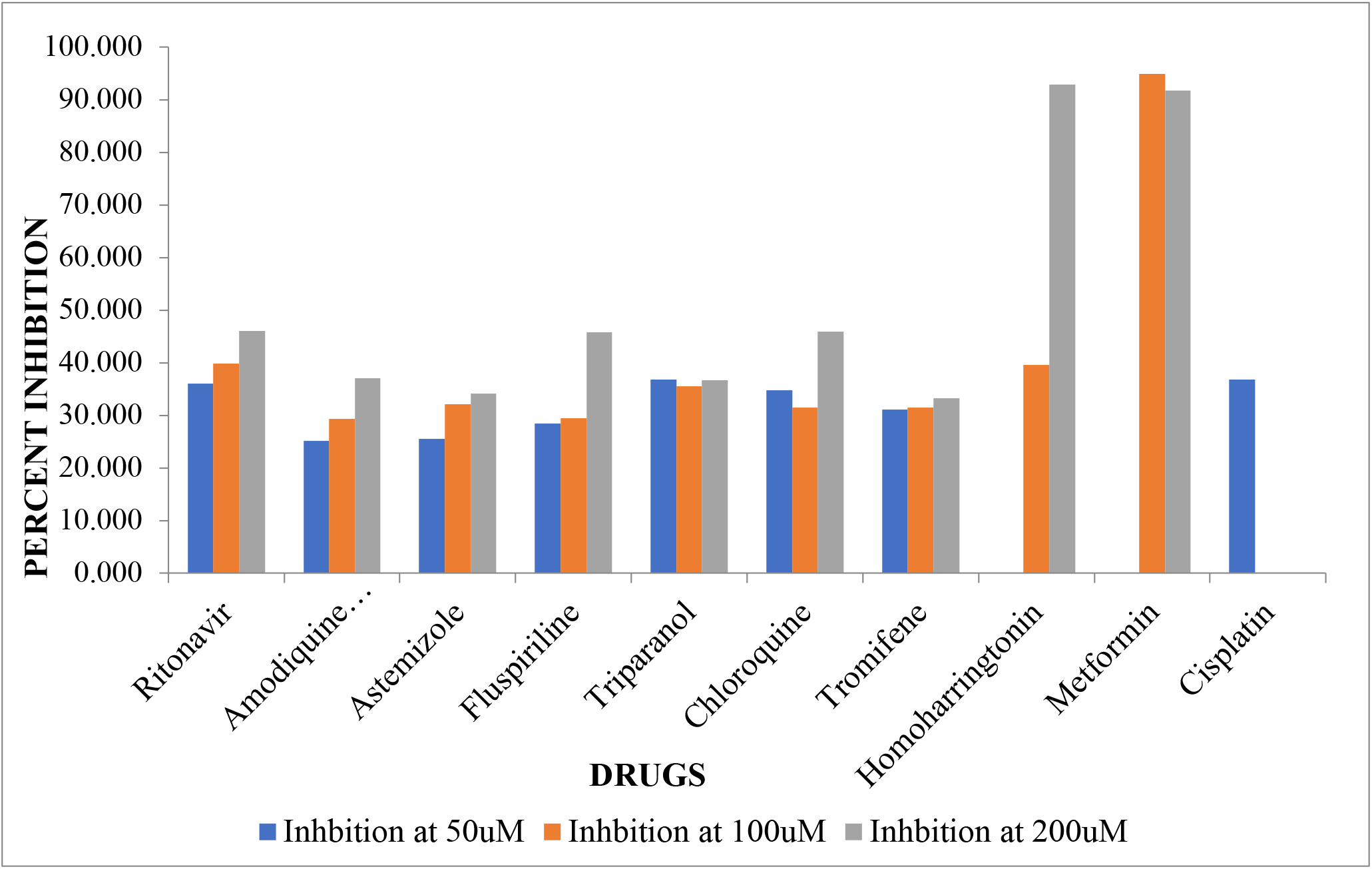
Inhibition of ACE2-RBD binding by drugs

### 2. Metformin prevents IL1b secretion

To establish whether metformin can block inflammatory stimulus (LPS) induced secretion of IL1b, a critical cytokine for propagation of COVI19 onset, we tested its effects in HUVEC cells in LPS treated cells. As can be seen in Figure 2, LPS stimulation increased secretion by close to 180 % of control. Interestingly only two of the drugs tested reduced the secretion of this cytokine, and metformin nearly reduced the secretion to control cell levels. This a direct demonstration that metformin is an anti-inflammatory drug which could have be very important in controlling inflammation in COVID19.

**Figure 2:**
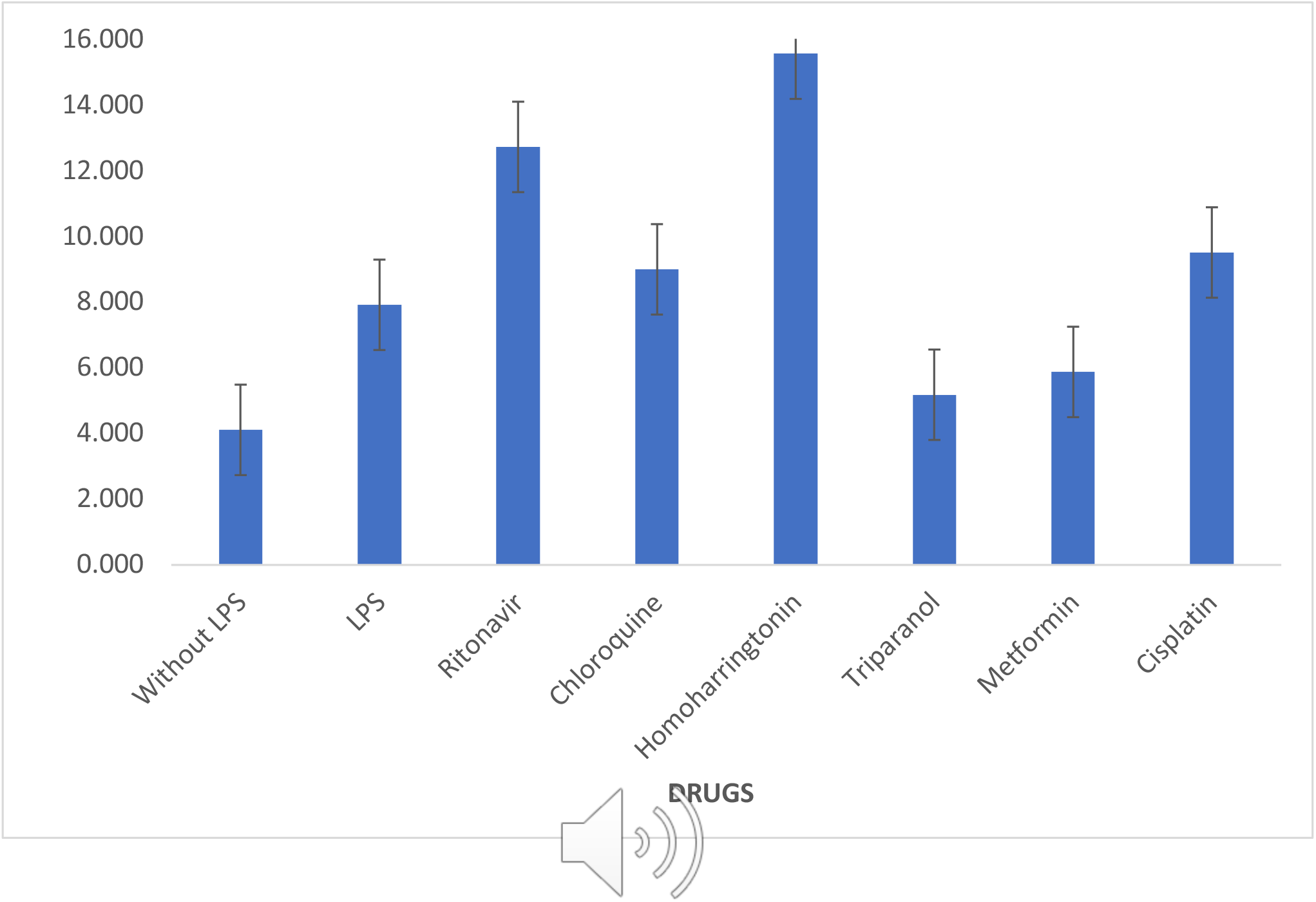
Secretion of ILb (femtogram/cell)-drugs added at 100uM

**Figure 3:**
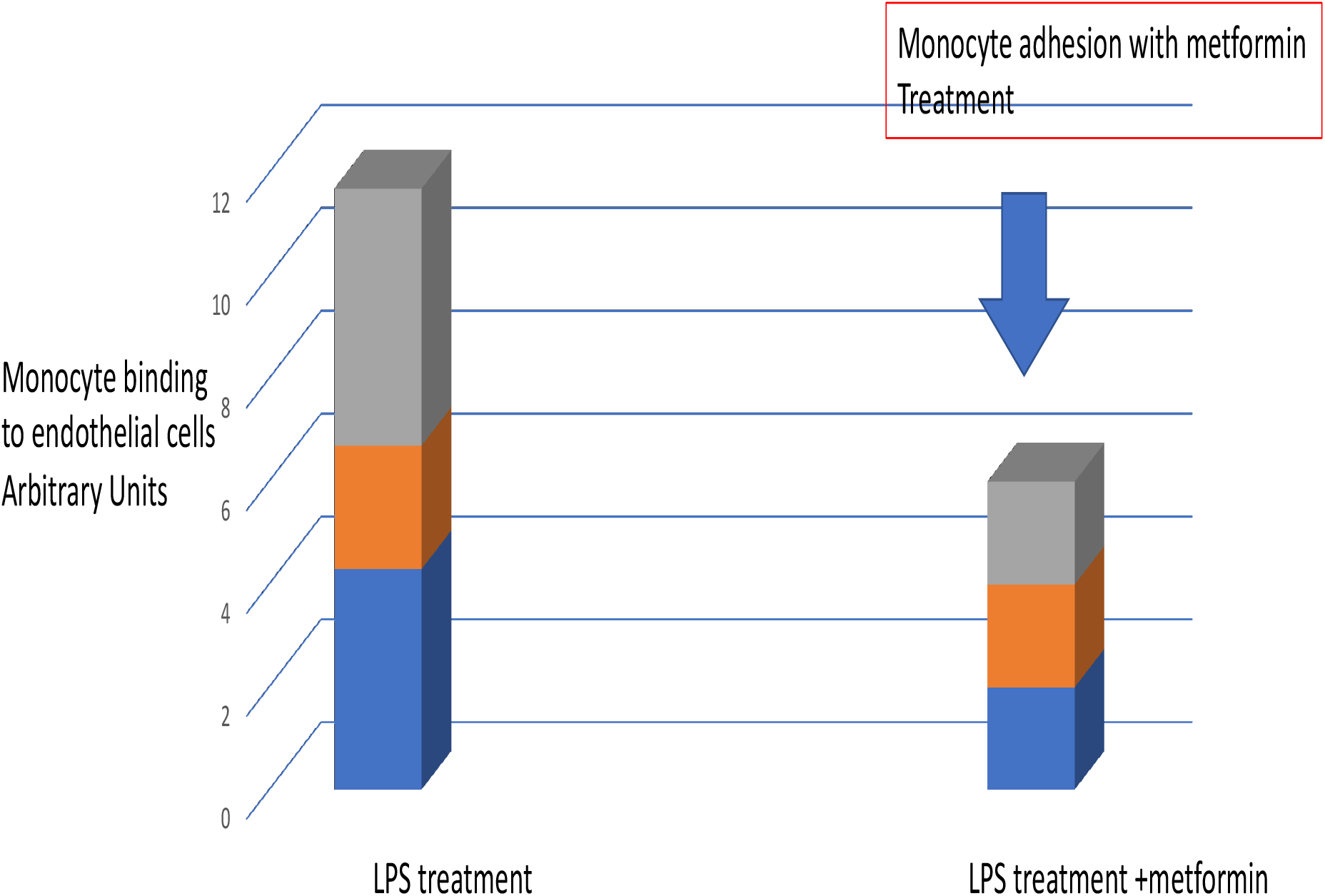
Inhibition of monocyte adhesion by metformin (200 uM)

### 3. Metformin reduces monocyte binding to endothelial cells

We next examined the ability of metformin to block the adhesion of monocytes to vascular endothelial cells. Adhesion of monocytes to inflammatory stimulus (LPS addition) activated cells is an necessary step for monocytes to enter the inflamed area. As shown below metformin at 200 uM treatment concentration reduced monocyte binding by more than 50% to LPS activated HUVEC cells. This reduction of monocyte accumulation by metformin is an indication that its anti-inflammatory activity extends beyond the cytokine secretion inhibition property shown above and may be beneficial in COVID19.

### 4. Metformin’s in vitro anti-infective activity and as nasal formulation in hamster model of COVID19

We then examined the effects of metformin on infection of SARS-CoV-2 virus in Vero E6 cells. In order to do this, the cytotoxicity of metformin in these cells at concentrations ranging from 0.25 to 3 mM if any, was first established. As shown in Figure 4, the drug was not cytotoxic as measured by LDH leakage assay when the cells were incubated with the drug. This suggests that the drug is not overtly cytotoxic to Vero E6 cells (maximum cell death was determined to be 7.4% at 3 mM).

**Figure 4:**
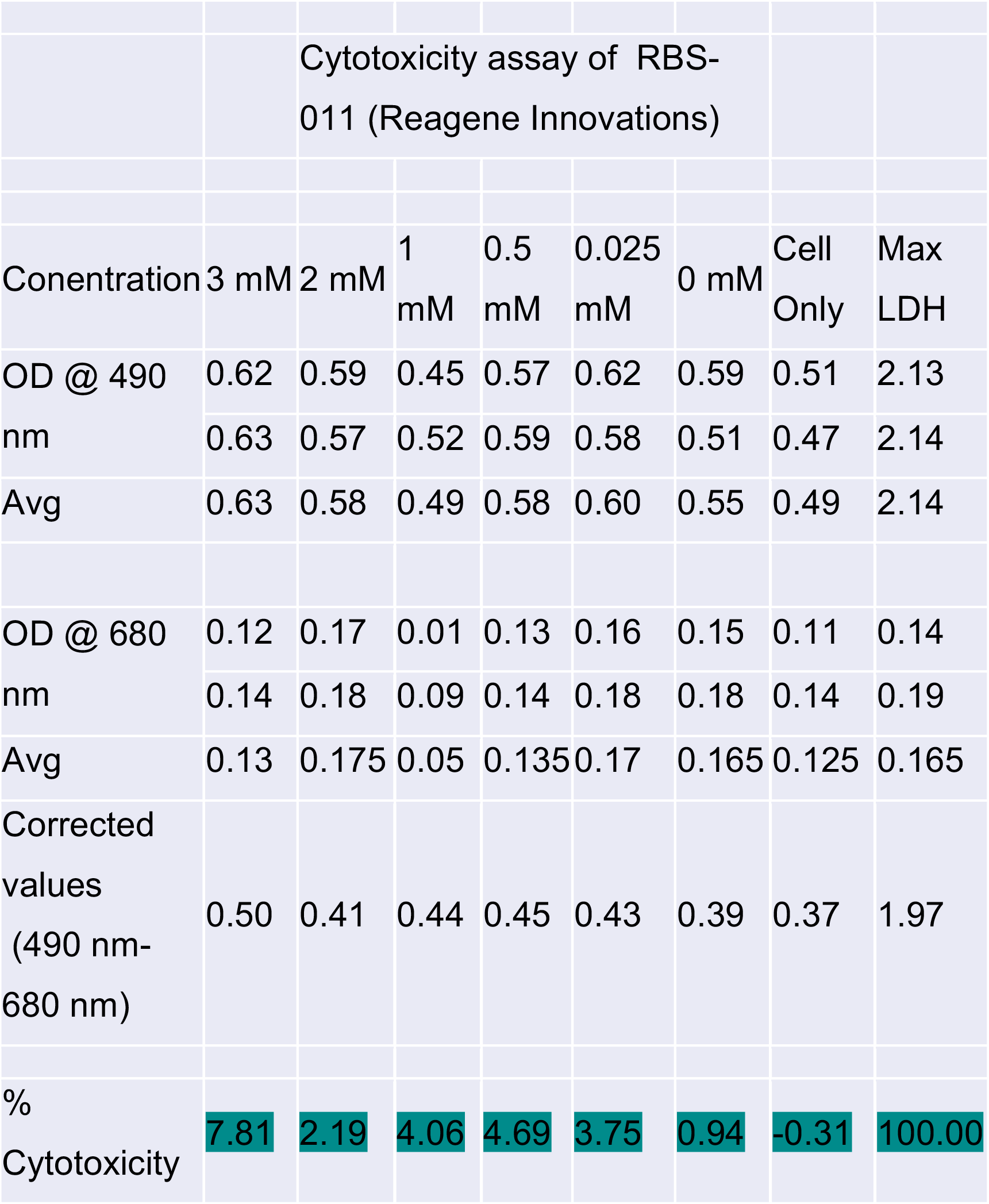
Cytotoxicity of metformin in Vero E6 cells

Next we examined the effect of metformin on infectivity of SARS-CoV-2 live virus in Vero E6 cells. As shown below in Figure 5 we tested remdesivir (as positive control) and metformin in the two intended formulations for in vivo study, i.e. deionized water and saline.

**Figure 5:**
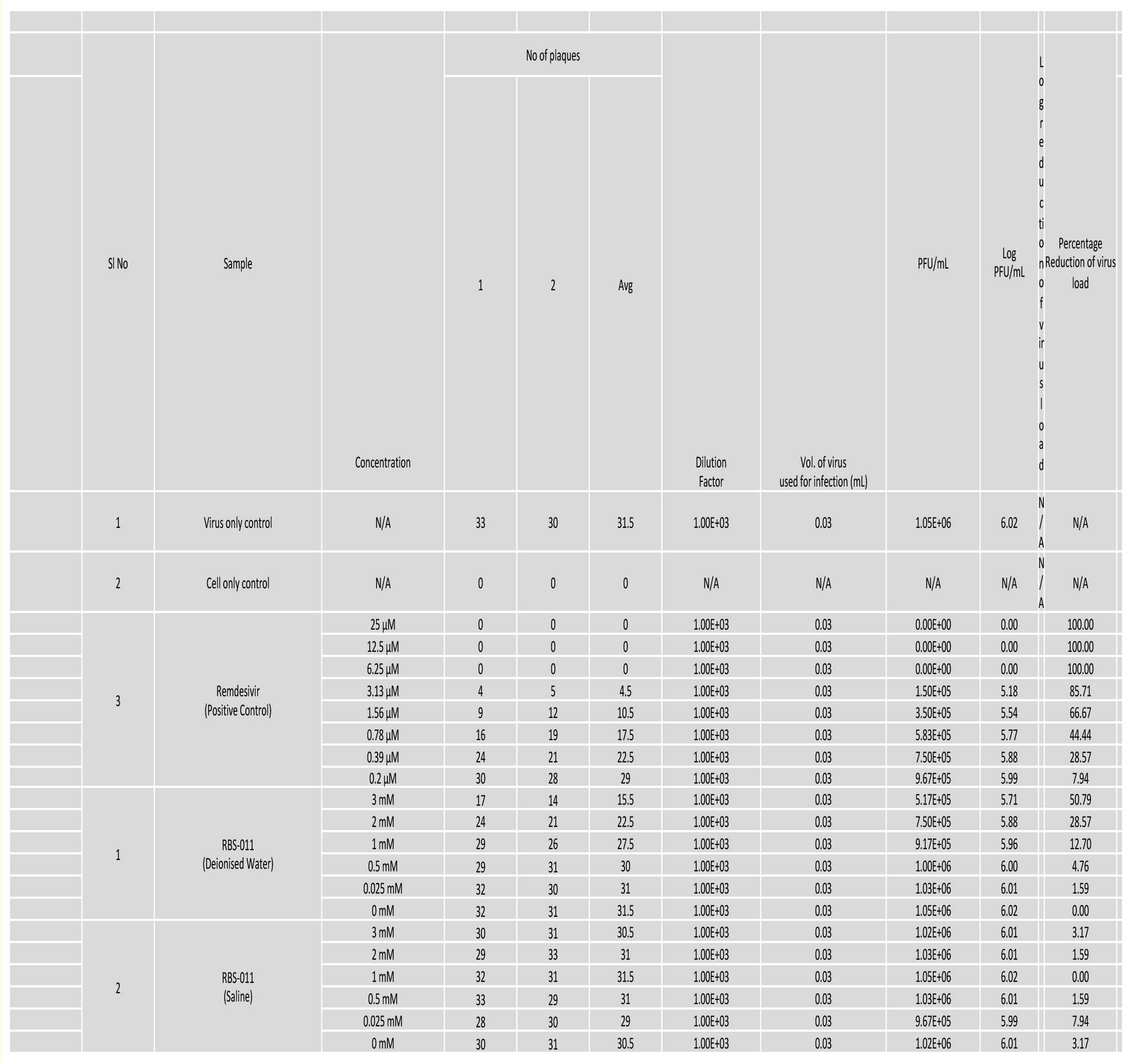
Metformin activity on blocking viral infection of Vero E6 cells

Remdesivir was effective in reducing viral infectivity with an IC50 of around 1.5 uM. Metformin in saline was not very active even at highest concentration tested (3 mM) but it reduced infection by 50% at the same concentration in deionized water. These data establish that metformin is active in blocking viral infection in deionized water. As such we used deionized water as the formulation for metformin in the in vivo studies.

#### Effect of metformin on SARS-CoV-2 infection in hamster model

Metformin has been proposed to be efficacious in COVID-19 in multiple preclinical and clinical studies (6,7,8, 9,10). But there are no reports of it being tested using a nasal route, which would be most relevant to COVI-19. So we tested Metformin, dosed through nasal intubation in the hamster SARS-CO-2 model of live virus infection. The summary of study design and results are shown in Figure 6. Remdesivir was used as a positive control drug. After 4 days of infection challenge, lungs were harvested and viral load determined. As shown below, remdesivir reduced viral load and this effect was statistically significant. Metformin at both doses tested (10 mg/ml and 50 mg/ml) had a more modest inhibitory effect as compared to remdesivir although there was reduction compared to untread control animals.

**Figure 6:**
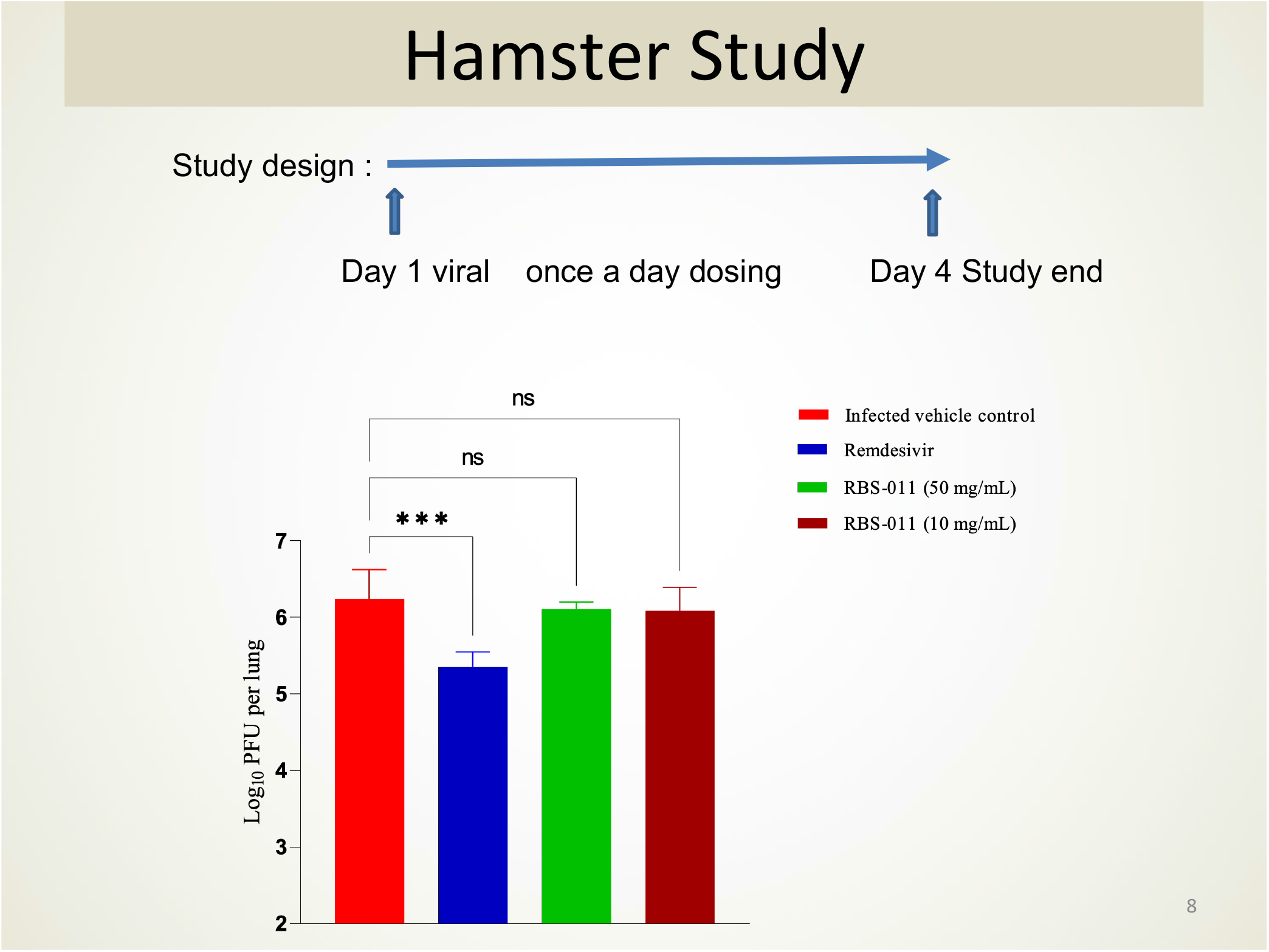
Effect of remdesivir and metformin in hamster model

## Discussion

We show here that metformin posses three important properties needed to have an effect on SARS-CoV-2 infection – it can block the entry of the virus into human cells thru the ACE2 receptors, it inhibits secretion of IL1b an important cytokine in the “cytokine storm” seen in COVID-19 and it inhibits the accumulation of inflammatory monocytes.

Metformin as a nasal formulation has never been tested before so, we used this route in our studies. While there was clear demonstration of the anti-inflammatory and anti-infective activity of metformin, its ability to reduce viral load in the lungs in the hamster model were modest compared to the positive control remdesivir. There could be several reasons for this observation

1. The dose we used (10 and 50 mg/ml) may not have been optimal and since the animals showed no adverse effects a higher dose can be attempted in future
2. The intra nasal delivery may not have delivered efficacious amounts of the drug to the lungs
3. The drug may have been metabolized/diluted rapidly in the lungs which is a heavily vascularized tissue
4. The dosing regimen using intra nasal delivery by intubation into the nares was tricky and may not have delivered the drug effectively

While these are some of the reasons which could be addressed in future studies, we believe our data have paved the way for metformin to be repurposed for COVID-19 after establishing optimal conditions of dose, delivery and frequency thru the nasal route. The robust anecdotal data in literature about the preclinical and clinical efficacy of metformin in COVID-19 strongly supports further development of a nasal formulation of this drug for COVD-19 (11,12,13,14,15,16). Since several of the steps observed in SARS-Co-V2 infection are common to many viral diseases, it may be useful to test this drug against other viruses as well.

## Acknowledgements

The authors are grateful for a BIRAC grant from Department of Biotechnology, India (DBT, India) to conduct some of these studies. We also acknowledge the support and advice of Dr. Sreedhar Narayanan and Mayas Singh at FNDR for the live virus studies.

